# Interoceptive signals shape the earliest markers and neural pathway to awareness at the visual threshold

**DOI:** 10.1101/2023.07.13.548857

**Authors:** Viviana Leupin, Juliane Britz

**Affiliations:** Department of Psychology, University of Fribourg, P.-A. Faucigny 2, CH-1700 Fribourg, Switzerland

**Keywords:** consciousness, brain-body interaction, EEG, cardiac phase, respiratory phase

## Abstract

Variations in interoceptive signals from the baroreceptors (BRs) across the cardiac and respiratory cycle can modulate cortical excitability and so affect awareness. It remains debated at what stages of processing they affect awareness-related event-related potentials (ERPs) in different sensory modalities. We investigated the influence of the cardiac (systole/diastole) and the respiratory (inhalation/exhalation) phase on awareness-related ERPs. Subjects discriminated visual threshold stimuli while their electroencephalogram (EEG), electrocardiogram (ECG) and respiration were simultaneously recorded. We compared ERPs and their intracranial generators for stimuli classified correctly with and without awareness as a function of the cardiac and respiratory phase. Cyclic variations of interoceptive signals from the baroreceptors (BRs) modulated both the earliest electrophysiological markers and the trajectory of brain activity when subjects became aware of the stimuli: an early sensory component (P1) was the earliest marker of awareness for low (diastole/inhalation) and a perceptual component (visual awareness negativity, VAN) for high (systole/exhalation) BR activity, indicating that BR signals interfere with the sensory processing of the visual input. Likewise, activity spread from the primary visceral cortex (posterior insula) to posterior parietal cortices during high and from associative interoceptive centers (anterior insula) to prefrontal cortex during low BR activity. Consciousness is thereby resolved in cognitive/associative regions when BR is low and in perceptual centers when it is high. Our results suggest that cyclic fluctuations of BR signaling affect both the earliest markers of awareness and the brain processes underlying conscious awareness.

**Significance Statement:** The brain continuously processes stimuli from inside and outside the body, and interoceptive stimuli can modulate the perception of external stimuli. Cardiac and respiratory rhythms are important pacemakers of the organism, and we show how they shape awareness-related brain activity for visual threshold stimuli in two ways. Variations of baroreceptor (BR) activity across the cardiac and respiratory cycle affect 1) the earliest electrophysiological marker (P1 for low (diastole/inhalation), VAN for high (systole/exhalation) BR activity) and 2) the brain areas activated (frontal cortex for low and parietal cortex for high BR activity) when subjects become aware of a stimulus. Cyclic variations of bodily signal can modulate cortical excitability and so shape the pathway to awareness and we propose to consider them as functionally relevant signals rather dismissing them as noise.

## Introduction

Trial-by-trial variations in brain activity modulate the response of the brain to external stimulation (1, 2) and can predict fluctuations in conscious awareness (3–5). Awareness of a stimulus requires the dynamic interplay of activity in sensory, parietal and frontal cortex (6–8). Both activity in a posterior occipito-temporo-parietal hot zone (9) and interactions between parietal and frontal cortex (6, 10) have been proposed to reflect awareness, but their respective roles remain debated. Frontal cortex is predominantly activated when subjects report their percept, but not in no-report paradigms (9), suggesting that frontal cortex activity reflects selective attention or response selection rather than awareness per se. It remains further debated whether the earliest electrophysiological markers of awareness reflect sensory (P1 (11)), or perceptual and post-perceptual (VAN, P3 (12, 13)) ERP components.

Awareness also consistently recruits the insula (14), a multi-modal hub with a caudo-rostral trajectory of connections to parietal and frontal areas (15–17). The Posterior Insular Cortex (PIC) is considered the primary visceral cortex (18) and processes interoceptive stimuli (19, 20); the Anterior Insular Cortex (AIC) integrates sensory stimuli both from outside and inside the body (17, 21) and together with the anterior cingulate cortex (ACC) forms the main hub of the saliency network (22, 23).

The involvement of the insula in awareness and the intricate connection between brain and body can explain why conscious awareness also varies with cyclic fluctuations in bodily signals. The cardiac rhythm is an important pacemaker of the organism, and increased blood pressure in the systole activates the baroreceptors (BRs), stretch sensors located in the aortic arch and carotid sinus which maintain a stable blood pressure through the baroreflex (24). BRs project via the nucleus tractus solitarii (NTS) to somatosensory cortex and the PIC (25, 26), and this ascending cascade of visceral afferents decreases cortical excitability (24), which can be conceptualized in terms of gain control. Gain is described by the sigmoid relationship between input and output strength, whose slope represents the differentiation between relevant (sensory) and irrelevant (interoceptive) signals. During high gain (silent BRs), relevant signals are amplified and irrelevant signal are attenuated (steep slope), whereas during low gain (active BRs), the differentiation between relevant and irrelevant signal is less pronounced (shallow slope) (27). Hence, BR activity can attenuate concurrent brain activity and so affect cortical excitability in a bottom-up fashion (24) which explains how it can modulate the processing of external stimuli: simple visual (28–30), auditory (31) and somatosensory (32–34) stimuli are detected more readily in the diastole when the BRs are inactive (27), and stimuli presented during the diastole elicit higher ERP amplitudes (32, 35).

Another rhythmic driver of the organism is respiration whose primary physiological function is gas exchange (O_2_ for CO_2_), and stimulus processing likewise varies across the respiratory phase: emotional (36, 37) and visual (37–39) stimuli are preferentially processed during inhalation, but somatosensory stimuli during exhalation (33). Subjects also synchronize breathing with task demands by inhaling around stimulus presentation (33, 38, 39) and exhaling when responding (39). Respiration can affect awareness in a direct and an indirect way: rhythmic mechanical stimulation of the olfactory bulb during inhalation directly drives the entrainment of rhythmic brain activity, and respiratory phase so modulates both oscillatory (36, 40–42) and broad-band resting-state (43) activity. The indirect influence of the respiratory phase on awareness is mediated by respiratory sinus arrythmia (RSA), the coupling of the cardiac frequency to the respiratory phase (acceleration during inhalation / deceleration during exhalation). RSA is a complex mechanism linking the cardiac and respiratory systems to dynamically adapt to changes in metabolic demands (44); it is partially modulated by the balance between the sympathetic and parasympathetic nervous system (45) and mediated by decreased BR activity during inhalation to optimize oxygen uptake (46, 47). Like for the cardiac phase, BR activity also fluctuates cyclically across the respiratory phase.

Despite their tight coupling, the influence of cardiac and respiratory phase on awareness-related brain activity have not been assessed jointly. We bridge this gap and investigate at what stages of processing the effects of cardiac and respiratory phase exert their influence on awareness-related ERPs in the visual modality. While interoceptive signals from the body clearly modulate the processing of sensory stimuli, the mechanism and timing underlying the interaction between interoceptive and sensory stimuli remains debated. An ERP study using somatosensory stimuli found reduced perceptual sensitivity in the systole and postulated that response to the predictable heartbeat is suppressed when the stimulus arrives with the cardiac pulse wave (32). It is unclear whether the heartbeat is suppressed in all sensory modalities or only if it shares the pathway with the sensory stimulus (48). The notion of cyclic fluctuations in gain control and cortical excitability as a function of BR activity (27) provides an alternative and more general framework for the interplay between interoceptive and sensory processing because it can explain the effects of BR activity fluctuations across both the cardiac and the respiratory phase. To differentiate between these two accounts on the influence of bodily signals on awareness, we investigate how and when cyclic variations of BR activity across the cardiac and respiratory phases affect ERPs elicited by visual threshold stimuli correctly classified with and without awareness (3, 49) (Fig. 1). Equating performance and physical stimulus properties avoids the confound of awareness and performance. We used ERPs and EEG Source Imaging (50, 51) to 1) contrast the time course of ERP amplitudes and their intracranial source differences elicited by aware and unaware identification in phases of high and low BR activity (low and high gain) across both the cardiac and respiratory cycle and 2) to show how cyclic variations of BR activity affect the interplay between sensory and higher cortical areas when subjects become aware of a visual stimulus.

**Fig 1.**
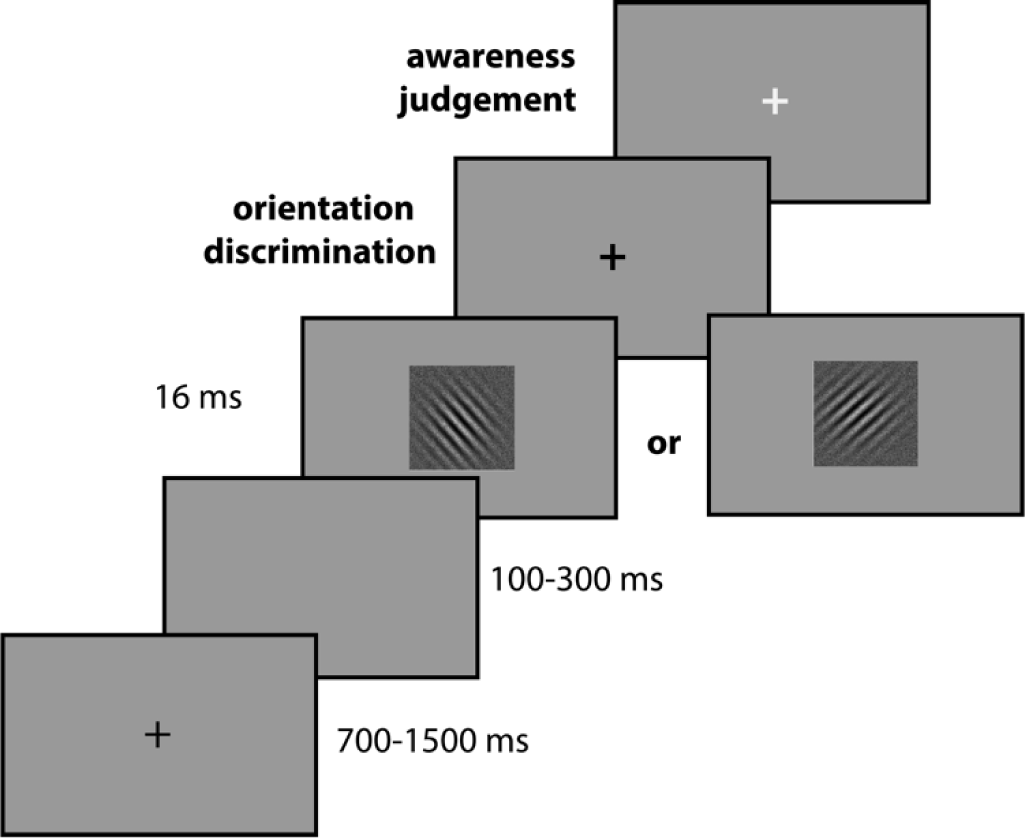
Experimental procedure. A Gabor grating either oriented to the left or to the right was presented for 16 ms. Subjects first reported the orientation of the stimulus (objective measure of accuracy) and then whether they saw the stimulus or whether they guessed (subjective measure of awareness).

## Results

### Behavioral results

Subjects responded correctly in 85.5% (SD=6.4%) of trials with a mean ratio of correct aware/unaware responses of 50.4/49.6% (SD=11.5%); only correct responses were submitted to further analyses. Subjects responded on average faster in the aware (755±196 ms) than the unaware (998±352 ms) condition (*SI Appendix*, Fig. S1). The GLMM revealed a main effect of awareness (estimate=114.75, SE=6.3, p=10^-16^) but no main effect of cardiac (estimate=0.0024, SE=3.09, p=0.99) or respiratory (estimate=0.89, SE=2.95, p=0.76) phase, and neither phase interacted with awareness (cardiac x awareness: estimate=-1.73, SE=4.54, p=0.7; respiratory x awareness: estimate=-0.023, SE=4.28, p=0.996).

### Stimulus-evoked potentials

On average, 251 trials were retained in the aware (126/125 (systole/diastole), 114/137 (inhalation/exhalation)) and 244 in the unaware (120/123 (systole/diastole), 112/132 (inhalation/exhalation)) condition in each subject. Despite being presented for only 16 ms, the stimulus elicited clearly discernible canonical ERPs.

### Awareness modulates ERPs

These ERPs were significantly modulated by awareness in three time-windows reflecting early sensory (P1, 90-120ms), perceptual (VAN, 250-350ms) and post perceptual (P3/LPC, 400-600ms) processes (*SI Appendix,* Fig. S2). We applied the False-Discovery Rate (FDR) (52) across all 150 time-points / 128 electrodes to correct for multiple comparisons. The P1 was more positive over occipital electrodes in the aware than the unaware condition; the VAN encompasses the N2 complex and was more negative in the aware than the unaware condition over occipital and lateral temporal electrodes, and the P3/LPC component was more positive in the aware than the unaware condition over centro-parietal electrodes.

### Cardiac phase modulates ERPs

The cardiac phase modulated visual ERPs (*SI Appendix,* Fig. S3) over occipital and lateral temporal electrodes in two time-windows: ERPs were significantly more positive in the systole than the diastole in the time-window -100-200ms and more negative in the time-window 230-550ms (*SI Appendix,* Fig. S3*A*). In the systole, the cardiac pulse wave overlaps with stimuli presented within this interval (-100-200ms); in the diastole, the same cardiac pulse wave appears to occur later because it is elicited during the systole of the subsequent heartbeat (300-500ms) (*SI Appendix,* Fig. S3*C*). This effect is not caused by the cardiac field artefact (CFA), the electrical field generated by cardiac activity that can potentially interfere with the EEG (53), (*SI Appendix,* Fig. S3*B*).

### Cardiac phase selectively modulates awareness-related ERPs

The cardiac phase differentially modulated awareness-related ERPs reflecting sensory (P1) and perceptual processes (VAN) (Fig. 2*C*). The amplitude of the P1 peak was selectively modulated by awareness: only in the diastole, it was attenuated in the unaware relative to the aware condition (cardiac phase x awareness: (F(1,29)=6.74, p=0.015, η^2^ =0.19, Fig. 2*B*). Linear contrasts revealed a significant effect of awareness in the diastole (*t*(*29*)=3.16, p=0.007, *d*=0.57), but not the systole (*t*<1) (Fig. 2*A*). Awareness and cardiac phase did not interact during the VAN time-window, but the effect size for the contrast between the aware and unaware conditions was larger in the systole than the diastole in the time-window 220-350 ms (Fig. 2*D*). The earliest marker of awareness varied with BR activity across the cardiac phase: P1 (inactive BRs) in the diastole and VAN (active BRs) in the diastole.

**Fig 2.**
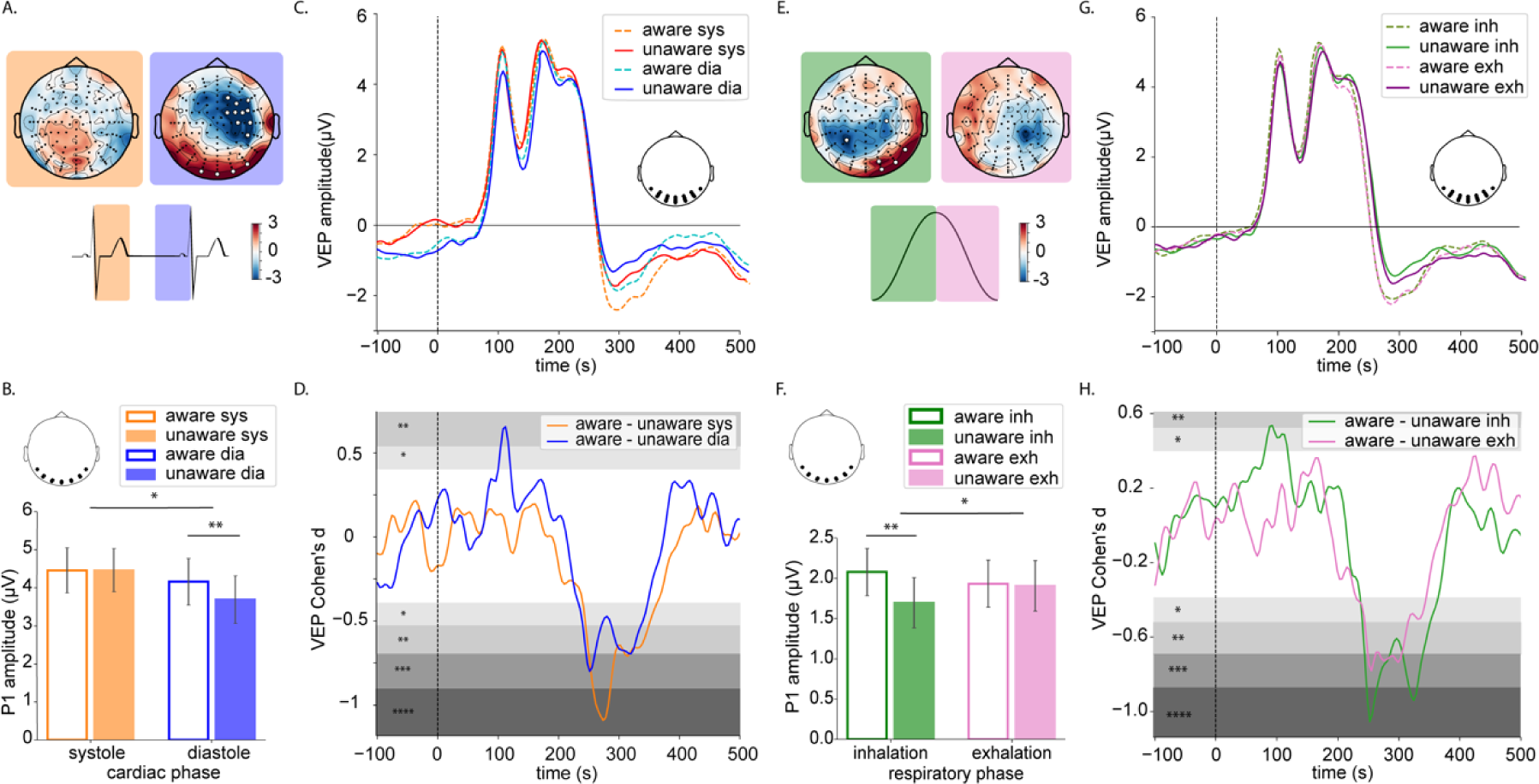
Modulation of the P1 and of the VAN as a function of awareness and cardiac and respiratory phase. (A) Topoplots of t-values for the contrast between aware and unaware conditions in the P1 time-window (90-120ms) in the systole (orange) and diastole (blue). Electrodes showing a significant interaction between awareness and cardiac phase are highlighted by white dots (FDR-corrected across all 150 timepoints and all 128 electrodes). (B) Significant interaction between awareness and cardiac phase on the P1 amplitude (90-120ms) averaged across occipital electrodes highlighted in the inset; error bars represent the standard error; *p<0.05, **p<0.01. (C) Grand Average ERP waveforms in all four conditions averaged across the electrodes showing a main effect of awareness in the time-window of the VAN (250-350ms) displayed in the inset. (D) Time-course of Cohen’s d effect sizes for the contrast aware - unaware in the systole (orange) and diastole (blue) at the electrodes highlighted in the inset in panel (C); gray shades indicate significance levels: *p<0.05, * p<0.01, ***p<0.001, ****p< 0.0001. (E) Topoplots of t-values for the contrast between aware and unaware conditions in the P1 time-window (65-95ms) during inhalation (green) and exhalation (pink). Electrodes showing a significant interaction between awareness and respiratory phase are highlighted by white dots (FDR-corrected over the entire time-window (-100-500 ms) and all 128 electrodes). (F) Significant interaction between awareness and respiratory phase on the P1 amplitude (65-95ms) averaged across occipital electrodes highlighted in the inset; error bars represent the standard error; *p<0.05, **p<0.01. (G) ERP waveforms in all four conditions averaged across the electrodes showing a main effect of awareness in the time-window of the VAN (250-350 ms) displayed in the inset. (H) Time-course of Cohen’s d effect sizes for the contrast aware - unaware during inhalation (green) and exhalation (pink) at the electrodes highlighted in the inset in panel (G); gray shades indicate significance levels: *p<0.05, **p<0.01, ***p<0.001, ****p<0.0001

### Respiratory phase selectively modulates awareness-related ERPs

Unlike the cardiac phase, the respiratory phase did not modulate ERPs. Like the cardiac phase, the respiratory phase selectively modulated awareness-related ERPs, albeit somewhat differently (Fig. 2*G*). The amplitude of the ascending slope of the P1 (65-95ms) was selectively modulated by awareness (respiratory phase x awareness: F(1,29)=4.65, *p*=0.039, η^2^ =0.14 Fig. 2*F*) and attenuated in the unaware condition only during inhalation (*t*(*29*)=3.32, *p*=0.004, *d*=0.6) but not exhalation (*t<1*) (Fig. 2*E*). Like for the cardiac phase, awareness and respiratory phase did not interact during the VAN (220-350 ms), but the effect size for the contrast between the aware and unaware conditions were larger during inhalation than exhalation (Fig. 2*H*). Unlike the cardiac phase that differentially affected the P1 and the VAN, the respiratory phase affected them similarly: both were more strongly pronounced during inhalation than exhalation. However, like for the cardiac phase, the earliest marker of awareness varied with BR activity across the respiratory phase: P1 during inhalation (inactive BRs), and VAN during exhalation (active BRs).

### Stimulus-evoked sources

We determined the sources underlying the sensor space effects by computing distributed inverse solutions and then applied statistical parametric mapping to compare the intracranial generators in the aware and unaware conditions during the time-windows of significant amplitude differences of the P1, VAN and P3/LPC.

### Cardiac phase modulates source activity underlying sensory processing

Unlike in sensor space, there was no main effect of awareness and no interaction between awareness and cardiac phase in the source space in the time-window of the P1 (90-120 ms). However, during that time-window, the cardiac phase differentially modulated source activity: current density was higher in the diastole than the systole in the aware (*t*(29)=3.45, *p*=0.003, *d*=0.63) but not in the unaware condition (*t*(29)=1.16, *p*>.05) (*SI Appendix,* Fig. S4*B*) in a spatio-temporal cluster encompassing right extrastriate visual and entorhinal cortex (MNI(max): x=34.3, y=-73.5, z=-12, *p*=.0272, *SI Appendix,* Fig. S4*A*).

### Cardiac and respiratory phase modulate awareness—related source activity underlying perceptual processes

Fig. 3*A-C*) summarizes the results of the source space analyses in the time-window of the VAN (240-340ms), all results are corrected for multiple comparisons across time and space using cluster-based permutation (54). During the entire VAN time-window, current density was higher in the aware than the unaware condition only in the systole in a spatio-temporal cluster encompassing the PIC and its ventral projection targets in the left temporo-parietal junction (TPJ): Superior Temporal Sulcus (STS) and Angular Gyrus (AG)); p=0.036, MNI(max): x=- 40.2, y=- 60.1, z=- 10.6, Fig. 3*A*), also note larger effect size for the main effect of awareness in sensor space in the systole than diastole.

**Fig 3.**
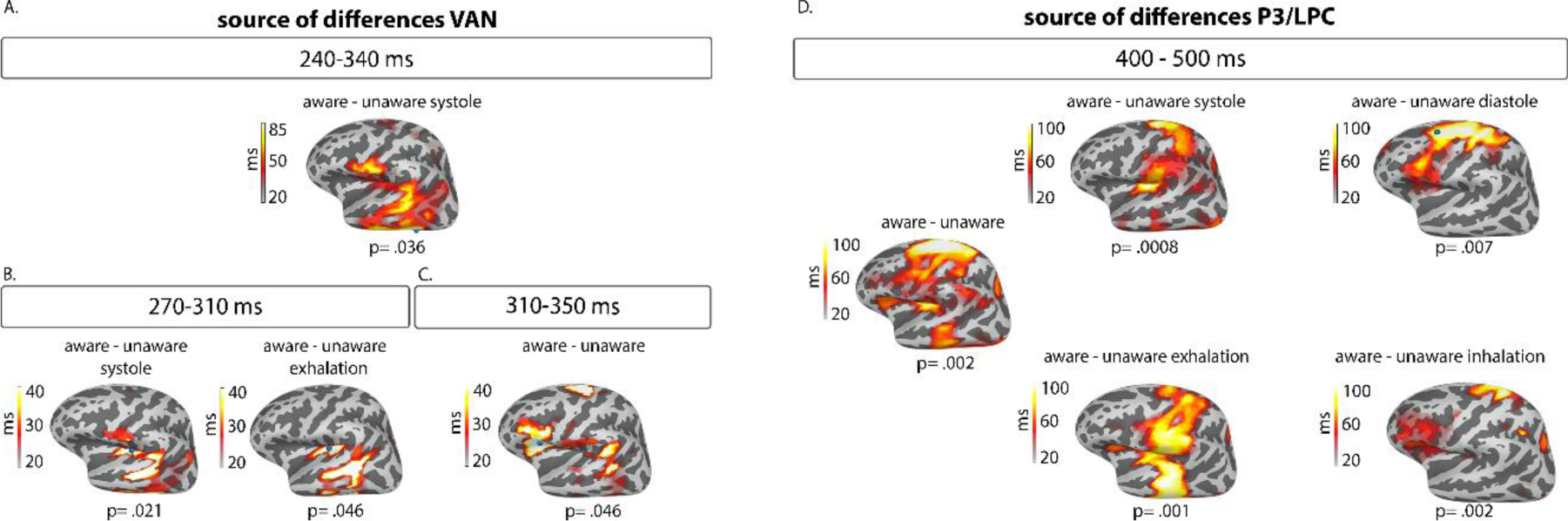
Current density differences underlying the main effect of awareness as a function of the cardiac and respiratory phase for the VAN and P3/LPC components rendered on the FsAverage surface template; color bars indicate the duration (>20 ms) of significant activity in the spatio-temporal clusters between the aware and unaware conditions at the respective significance level. (A) Main effect of awareness in the systole during VAN time-window (240-340 ms). (B) Main effect of awareness during the early VAN time window (rising phase, 270-310 ms in the systole (left) and exhalation (right). (C) Main effect of awareness in the late VAN time window (310-350 ms) across all conditions. (D) Main effect of awareness during the entire LPC window (400-500 ms) across all conditions (left), for the systole (top left), diastole (top right), exhalation (bottom left) and inhalation (bottom right).

Because activity differences persisted for a long time (100ms), we investigated the temporal evolution of source activity differences and sub-divided the VAN time-window into an early (270-310 ms) and a late (310-350 ms) part corresponding to the rising and falling phases of the VAN. In the early time-window, current densities were higher in the aware than the unaware condition in PIC and its ventral projection targets in STS only when BR activity was high (systole: p=0.021, MNI(max): x=-32.4, y=-31.9, z=17.1, exhalation: p=.046, MNI(max): x=-32.4, y=-31.9, z=17.1 (Fig. 3*B*)). In the late time-window, current densities were significantly higher in the aware than the unaware condition irrespective of the cardiac or respiratory phase in a cluster centered around AIC and Inferior Frontal Gyrus (IFG) that also encompassed the ventral section of the TPJ (STS, AG), (p=.046, MNI(max): x=-37.6, y=15.5, z=9.8, Fig. 3*C*).

### Cardiac and respiratory phase modulate awareness—related source activity underlying post-perceptual processes

Fig. 3*D* summarizes the results of the source space analyses in the P3/LPC time-window (400-500 ms). Current densities were higher in the aware than the unaware condition irrespective of cardiac or respiratory phase both in the PIC and AIC and their respective dorsal projection targets in parietal (TPJ(SMG), IPL) and frontal cortices (IFG, SFG), along with medial areas extending posteriorly (Precuneus, Posterior (PCC) and Middle (MCC) Cingulate and Fusiform Gyrus (FFG)) and anteriorly (ACC, SFG); p=0.002, MNI(max): x=-11.5, y=-7.7, z=47.3 *SI Appendix,* Fig 5). Medial sources did not differ between conditions, but like during the VAN time-window, lateral sources differed along a posterior-anterior gradient that varied with cyclic variations of BR activity: during high BR activity, PIC is reactivated along with its dorsal projection targets in SMG and IPL in the systole (p< 0.001, (MNI(max): x=-4.1, y=-33.4, z=30.5)); AIC and middle temporal gyrus (MTG) are additionally activated during exhalation (p=0.001, (MNI(max): x=-4.1, y=-33.4, z=30.5)). During low BR activity, AIC is re-activated along with its projection targets in IFG and ACC (inhalation, p= 0.002, (MNI(max): x=-4.1, y=-33.4, z=30.5); SFG is additionally activated during diastole (p= 0.007, (MNI(max): x=-30.8, y=9.3, z=53.6).

## Discussion

We show that cyclic variations of interoceptive signals across the cardiac and the respiratory phase determine both the earliest electrophysiological markers of awareness and the trajectory of brain activity when subjects become aware of a visual threshold stimulus.

The P1 was the earliest ERP component modulated by awareness when BR activity was low, and gain was high (diastole, inhalation). Current density in extrastriate visual cortex was lower in the systole than the diastole when subjects saw the stimulus, which indicates that BR activity can reduce cortical excitability in response to a near-threshold stimuli when it is detected. The modulation of the P1 with cyclic fluctuations in BR input helps to solve the controversy whether it reflects the earliest marker of awareness (11) or not (55): it does but only when concomitant interoceptive input from the BRs is minimal.

The VAN was the earliest marker of awareness when BR was high, and gain was low (systole, exhalation). It was significantly modulated by awareness irrespective of the cardiac or respiratory phase, however, the effect sizes of the contrast between the aware and unaware conditions were larger in phases of high BR activity / low gain (systole, exhalation), indicating an increased processing demand when the visual stimulus occurs along with signals from the BRs and both stimuli have to be processed. Its intracranial generators differed both across time and with cyclic variations in BR activity along a posterior to anterior gradient of spread of activation from the insula. The PIC and its projection targets in the ventral TPJ and was exclusively activated during the early (rising) phase of the VAN and only when BR activity was high (systole/exhalation). The PIC is considered the primary interoceptive cortex (18) and is the main cortical projection target of the BRs (25, 26), and the TPJ receives and integrates multimodal inputs (56) and is considered a hub important for conscious awareness (57). When a visual stimulus is presented shortly after the heartbeat or during exhalation when the BRs are active, both visceral cortex and multimodal integration areas are activated jointly to effectively process and separate the visceral and visual inputs. This finding is in line with the notion of decreased gain (poorer separation between relevant (visual) and irrelevant (visceral) input) when the BRs are active (27). Our data suggest that this separation likely occurs in the TPJ, which is more strongly activated when subjects perceive than miss the stimulus. In other words, only when this separation between the visual and visceral stimuli is successful, subjects will perceive the visual stimulus, i.e. low gain (high BR activity) does not per se preclude awareness of a stimulus, but it requires more effortful separation between relevant and irrelevant inputs. No such separation of inputs is necessary during high gain / silent BRs and hence there is no difference in brain activity in this time window during diastole and inhalation. We show statistically significant differences in activity between aware and unaware identification of a visual stimulus in temporal cortex in this early time-window when subjects report their percept. The global neuronal workspace (GNW) model postulates that conscious awareness can only arise if activity reaches frontal and parietal areas after ∼300 ms, it also explicitly states that activity in occipital and inferior temporal areas before ∼300 ms does not differ between aware and unaware perception (6, 58). These areas are activated as a function of awareness earlier than postulated by the GNW model but only when the visual stimulus is accompanied by a visceral input, i.e. when gain is low and the brain has to process and separate the visual and visceral inputs. When this separation does not take place, subjects miss the stimulus. The increased TPJ activity is not driven by the visual stimulus when it reaches awareness per se, but it arises from increased processing demands when a visual and a visceral stimulus occur simultaneously.

More anterior areas (AIC, IFG) are activated along with the TPJ in the late (falling) phase of the VAN irrespective of the cardiac or respiratory phase. The AIC integrates visceral inputs with sensory, emotional, social and cognitive information relayed from other cortical areas (59). The AIC and IFG are important hubs of the SN (22, 23), which amplifies the salience of stimuli. Our findings suggest that 1) the VAN component reflects the critical transition at which salient inputs are selected and successively spread and maintained in a global conscious state, and 2) this process differs with cyclic variations in bodily signals from the BRs, which can add to our understanding of the role of the insula for awareness (14).

The source differences of the P3/LPC corroborate how cyclic variations in visceral input influence which brain areas are activated when subjects become aware of a visual stimulus. Overall, activity was stronger in widespread areas including the insula (AIC, PIC) and their respective dorsal and ventral projection targets along with medial areas when subjects became aware of the stimulus. However, cyclic variations of BR activity across the cardiac and respiratory phase recurrently and selectively reactivated the PIC when BR activity was high (systole/exhalation) and the AIC when BR activity was low (diastole/inhalation), and activity spread dorsally to parietal (from PIC) and frontal (from AIC) cortex when subjects become aware of the stimulus. This illustrates the modulatory role of primary visceral (PIC) cortex for awareness: it is activated both by phasic (systole) and tonic (exhalation) BR activation, and its role in conscious awareness is not limited to processing visceral signals when they co-occur with an external stimulus. Rather, the configuration of activity in the insula during the VAN determines the configuration of activity during the P3/LPC: awareness is supported by parietal cortex for high and by frontal cortex for low BR activity. Thus, none of these areas seem to be per se necessary for awareness but they can contribute independently to the awareness of stimulus.

Our findings shed a new light on the debate about the roles of parietal and frontal cortex for awareness: previous studies using no-report paradigms suggest that activity in the posterior parietal hot zone reflects awareness (9) and that PFC reflects selective attention and response selection rather awareness per se when subjects overtly report their percept (6, 10). We provide a new solution to this debate and show that parietal and frontal areas are differentially recruited as a function of both phasic and tonic cyclic variations in BR activity across the cardiac and respiratory phases rather than by task demands. The activity in more domain-general areas in posterior medial cortex (PC/PCC) does not vary with fluctuations in bodily signals. Taken together, variations in BR activity influence which sub-region of the insular cortex is recruited and hence to which cluster of connected regions the activation spreads. High BR firing activates the PIC in the early stage of the VAN which drives the signal to reverberate in posterior and highly connected parietal regions later on (similar to the posterior “hot zone”(9)). In phases of weaker BR activity stimulus processing is driven by the AIC-IFG cluster in the later stage of the VAN from where it spreads to high-order frontal cortices (in line with GNW predictions).

The interaction between awareness and cardiac and respiratory phase is not reflected in the reaction times. Subjects do respond faster when they perceive than miss the stimulus, but not differentially across the cardiac and respiratory phases. This is most likely due to our experimental paradigm: we compared only correctly classified stimuli with and without awareness, i.e. we did not include any errors to avoid having erroneous hits. Other experiments that separately considered hits vs. misses and correct vs. error trials (32) found more hits in the diastole than systole and more errors in the systole than diastole.

Our findings suggest that visceral signals are processed and separated from the visual input rather than suppressed (32), at least when visceral and sensory signals do not share the same neuronal pathways. The heart is the first organ that develops and starts to beat rhythmically at four weeks of gestation (60). The cardiac rhythm can be considered as the first pacemaker of a mammalian organism and is fully developed by the time the neural tube starts to form (61), and we show how signals from the heart shape the neural pathway to awareness. Respiration is another fundamental bodily rhythm whose physiological function goes beyond gas exchange; it can modulate oscillatory (36, 40, 41) and broad band (43) brain activity and modulate ERPs in a visual-spatial task (38), and we show here that cyclic fluctuations of BR activity across can likewise modulate awareness-related ERPs and their sources. Subjects breathed nasally, thus respiration can affect awareness both directly through entrainment of brain activity by means of OB stimulation (36, 42) and indirectly through RSA (46). Future studies using oral respiration will be needed to differentiate between the relative contributions of these routes.

The influence of cardiac and respiratory activity on the EEG is generally dismissed as noise and eliminated (62, 63). Our demonstration of how cyclic variations of bodily signals from the BRs across the cardiac and respiratory phase cyclically modulate cortical excitability and so shape the neural pathway to awareness. They modulate brain activity and affect awareness similar to trial-by-trial fluctuations in brain activity that are no longer eliminated as noise but considered to carry functional relevance. We propose to consider ECG and respiration as functionally relevant signals and to record them activity along with EEG as potentially important modulators of brain activity to open up new avenues to elucidate the intricate brain-body-connection.

## Materials and Methods

### Participants

Forty healthy right-handed subjects (26 female, age: 24.6 ± 5 years, range 18-42) participated in the study. None reported a history of neurological, psychiatric, cardiological and respiratory impairments and all were right-handed (64). The data from ten subjects were not retained for further analysis (six subjects did not meet the behavioral criteria of >75% correct answers and roughly 50/50% CA/CU for each stimulus orientation (one subject was excluded after the pre-test, in two subjects the EEG session was prematurely terminated, and data from three subjects were discarded after the EEG experiment), and four had compromised electrophysiological data quality (three showed abnormal ECG T-waves, and one had noisy EEG data). The data of 30 subjects (17 female, age 25.23±5.42 years, range 18-42) was submitted to further analysis. Participants received monetary compensation (20 CHF/hour) or course credits after given written informed consent; the full study protocol was approved by the Ethics Committee of the University of Fribourg.

### Stimuli and procedure

Stimuli were Gabor gratings subtending a visual angle of 5° (3 cpd) oriented left (135°) or right (45°) embedded in grayscale random dot noise. All stimuli were generated and presented using Psychopy3 (65) on a grey background on a ViewPixx Screen (1920 × 1080 pixel resolution, 120 Hz). Subjects were positioned on a chin rest 70 cm from the screen in a dimly lit room and performed the task while breathing through their nose; a piece of surgical tape was lightly placed over their mouth to prevent them from breathing orally.

First, a fixation cross was presented (700-1500 ms), followed by a blank screen for 100–300 ms and the target for 16 ms. Subjects first reported the perceived orientation by pressing the “F” (left index) or “J” (right index) key on a keyboard, (objective measure of accuracy) and then to indicate whether they saw (“J”) the stimulus or not (“F”) (subjective measure of awareness).

Prior to the EEG experiment, the perceptual threshold was titrated for both stimuli in each subject: the strength of the Gabor grating and the opacity of the random dot mask remained constant, and visibility was controlled by linearly varying the Michelson contrast of the random dot noise in 20 steps (66). Noise contrast levels that yielded an overall high accuracy (>75%) and roughly equal numbers of correct identification with and without awareness were retained for the EEG experiment. To control for threshold stability, the noise contrast could be adjusted if necessary, and 960 stimuli were presented in 12 blocks of 80 trials.

### Electrophysiological Recordings Data Preprocessing

EEG was recorded from 128 active Ag/AgCl electrodes (BioSemi®) referenced to the CMS-DRL ground. The electrocardiogram (ECG) and respiration were recorded as external bipolar channels using a bipolar montage (right clavicle/lower left rib) and a breathing belt (SleepSense®) placed on the lower abdomen. All physiological signals were simultaneously recorded at 1024Hz/16 bit.

Only correct responses with (aware) and without (unaware) awareness were analyzed. Trials were classified according to the cardiac (systole/diastole) and respiratory (inhalation/exhalation) phase that were identified using the Python Neurokit2 toolbox (67). Markers were placed on the R-peak and the end of the T-wave, denoting the beginning of systole and diastole. Because the diastole can be almost twice as long as the systole, we equalized their durations to have equal numbers of trials by only retaining stimuli falling within the interval at the end of the diastole corresponding to the duration of the systole in that cycle (32). Likewise, the peaks and troughs of the respiration trace were marked to denote the periods of inhalation and exhalation. Respiratory cycles 2.5 standard deviations faster or 1.5 standard deviations slower than the mean were excluded from further analyses.

All EEG analyses (pre-processing, averaging, source modeling and statistical analyses) were performed using the MNE-python toolbox version 0.24.0.1 (68). First, the EEG signal was algebraically re-referenced to the common average reference, filtered between 0.5 and 40 Hz (FIR filter, transition window of 10 Hz) and down-sampled to 256 Hz. Oculomotor and myogenic artifacts were removed using independent component analysis (ICA) (69). Trials contaminated by ocular artifacts occurring 300 ms before and after stimulus onset were rejected, all other oculomotor and myogenic artifacts were corrected by removing the respective ICs, and artifact-free components were forward-projected for further analysis. To investigate the potential contribution of the CFA, we separately created and analyzed a dataset retaining and excluding the CFA ICs. Next, the data was epoched between -200 ms to 1000 ms around stimulus onset, and artifact rejection and channel interpolation was performed in a data-driven way using the Autoreject procedure (70) implemented in MNE.

### Analysis of behavioral data

Only correct responses with and without awareness were analyzed, and outliers (reaction times (RTs) longer than the 97.5^th^ and shorter than the 2.5^th^ percentile) were omitted. Because RTs were not normally distributed, we used General Linear Mixed Effects Model with an inverse link (71) and treated subjects as a random factor to separately investigate the effects of awareness (aware/unaware), cardiac (systole/diastole) and respiratory (inhalation/exhalation) phase on the RTs.

### Analysis of stimulus-evoked potentials

The epoched data were averaged separately for each condition in each subject. We used mass univariate ANOVAs to assess statistical differences in ERP amplitudes as a function of the experimental factors awareness and cardiac or respiratory phase in the time-window -100-500ms around stimulus onset.

We made no a-priori assumptions for either localization or timing of effects and used FDR (52) to correct for multiple comparisons across time and space. Subsequently, we used the time-windows and electrode locations of the significant effects obtained in the ANOVAs as regions of interest (ROIs) for which we assessed the direction of significant differences between experimental conditions using mass-univariate paired-t-tests.

### Analysis of stimulus-evoked sources and source differences

The topography of the scalp electrical field represents the spatial summation of all currently active intracranial sources and serves as the precursor for source localization (*<u>SI Appendix,</u> <u>Methods</u>* for details). We computed the intracranial generators of the stimulus-evoked responses based on the scalp topography at each time point and for each experimental condition in each subject with the dSPM inverse solution (72) implemented in MNE. The source space was defined in a surface-based template of the cerebral cortex (FsAverage) (73) comprising 5124 solution points spaced equidistantly throughout the surface of the cortical gray matter including Insular Cortex. The forward model was solved using a realistic three-layer (brain, skull, scalp) boundary-element head model to construct the lead field. The spherical electrode coordinates provided by MNE were co-registered with the scalp surface, and finally, the inverse operator was applied to the evoked responses to obtain the source time course separately for each condition in each subject.

We restricted the statistical analysis of source differences to the time-windows corresponding to significant sensor-space differences (P1, VAN, P3/LPC). In the so-defined time-windows, we performed time-point-wise ANOVAs with the factors awareness and cardiac phase and awareness and respiratory phase, respectively. We used spatio-temporal cluster-based permutation tests to correct for multiple comparison correction across time and space (54, 74).

### Data and Code Availability

The consent forms signed by participants do not allow us to give free access to data but require us to check that data are shared with members of the scientific community. We can make the data available upon request to researchers.

## Acknowledgments

This research was funded by Swiss National Science Foundation grant 10001C_189408 to J.B. We thank Roberto Caldara for providing the lab infrastructure and Alen Jelusic, Dunja Vulliemin, Amira El Hachimi, Jade Ueberschaer, Samuel Müller, Fania Maffeis and David Elmiger for help with data collection.

