## Supplementary Material for "Interoceptive signals shape the earliest markers and neural pathway to awareness at the visual threshold"

* Juliane Britz

This PDF file includes:

Supplementary text
Figures S1 to S5
SI References

Supplementary Methods

#### *Determination of Discrimination Threshold and Awareness*

Subjective awareness at the sensory threshold is usually assessed by comparing hits vs. misses of the same stimulus irrespective of the objective performance in a given trial (1–4). Subjective awareness and objective performance are generally confounded (aware stimulus identification should yield errors, whereas unaware stimulus identification is as likely to produce errors and correct responses). However, awareness and performance are independent in the sense that awareness is not necessary for correct performance (5). To isolate the neural correlates of subjective awareness, we sought to avoid the confound between awareness and performance and kept performance constant for the same stimuli: we compared the same stimulus when it was classified correctly with (correct aware, CA) and without (correct unaware, CU) awareness (6), thereby excluding erroneous responses classified as aware (6). Standard adaptive staircase procedures (7) can only adapt one variable (e.g., accuracy > 80% or awareness 50%) at a time, but they cannot adapt two variables simultaneously (e.g., 50% awareness and > 80% accuracy). We aimed to have high accuracy and roughly equal numbers of aware and unaware trials and hence determined the discrimination threshold in a two-step behavioral pre-test. The procedure was identical to that in the main experiment (fixation cross for 700 – 1500 ms, blank screen for 100 – 300 ms, stimulus 16 ms, orientation discrimination ("F" for left, "J" for right) followed by awareness judgment ("F" for guessed, "J" for seen), see Figure 1).

Stimuli were Gabor gratings oriented left (135°) or right (45°) embedded in grayscale random dot noise. In a first step, we presented each subject with 20 stimuli for each orientation with decreasing visibility by increasing the Michelson contrast of the random dot noise mask from 30% to 100% in 20 steps to determine the approximate contrast level at which subjects stopped perceiving a stimulus. In case subjects perceived all / none of the stimuli, the opacity of the noise mask was decreased / increased to adjust visibility, and this step was repeated. In the second step, we redefined 20 stimuli for each orientation for which the Michelson contrast was centered around (+/- 20%) the value determined in the first step. Those stimuli were presented twice in a pseudo-randomized order in 5 experimental blocks (10 repetitions for each stimulus). This procedure took about 30 minutes to complete. For each orientation, we then selected the stimulus that had the highest accuracy (> 75%) and was closest to the desired 50/50 ratio of CA/CU trials for each orientation. This was done to avoid that performance was confounded with stimulus orientation to avoid that subjects consistently perceived one orientation but not the other. This procedure could dissociate objective performance and subjective awareness and so rule out stimulus conditions in which subjects responded erroneously despite reporting to have seen the stimulus, which indicates the subjective awareness judgments truly reflected conscious perception rather than overconfidence in stimulus classification.

The so-defined stimuli were used in the subsequent EEG experiment. Only data from those subjects who fulfilled the same behavioral criteria throughout the EEG experiment were retained for further analysis. Simultaneously controlling multiple parameters (high performance (> 75%) with roughly 50/50% CA/CU trials for each stimulus orientation) imposed strong constraints and could not be achieved in every individual throughout the study.

Of 40 subjects that initially participated, six subjects had to be excluded because of these strict behavioral criteria. One subject was excluded after the pre-test (low accuracy in the CA condition) and did not participate in the EEG experiment. Out of the 39 subjects who participated in the EEG experiment, in one the behavioral criteria could not be achieved, and the EEG session was terminated prematurely. Four subjects were excluded after completing the EEG session for not fulfilling the behavioral criteria (skewed ratio (20% CA/80% CU) for the right-oriented stimulus, overall low accuracy (68% and 56%), and low accuracy in the CA condition in the second half of the recording). Of the remaining 34 subjects, three were rejected due to abnormal T-waves in the ECG one due to compromised EEG signal quality.

*Reconstruction of EEG Sources and Inverse Space Statistics*

#### Creation of inverse operator and reconstruction of intracranial sources

The topography of the scalp potential field represents the spatial summation of all concurrently active intracranial current sources and serves as the precursor for the localization of its intracranial generators (inverse solution). However, any given topography can in principle be generated by an infinite combination of intracranial generators (8) that can vary in location, strength, extent and orientation, which is why the so-called inverse-problem is ill-posed and intracranial generators cannot be derived from the configuration of the scalp electrical field alone. To overcome this ill-posed inverse problem, a forward solution is required which incorporates biophysical constraints on source generation and propagation. This forward model comprises a source model that restricts the potential loci of intracranial currents and a lead field that 1) models the propagation of intracranial currents to the surface of the scalp and 2) accounts for the different conductivity values of the brain, skull and scalp resulting in blurring of the sources according to which the inverse problem can be solved using Maxwell-Equations (9).

Because the EEG is generated by postsynaptic dendritic currents in pyramidal cells (10), the source space has to be restricted to the cortical gray matter, and because all neurons are aligned perpendicularly to the cortical sheet, so is the orientation of the local intracranial generators. We used the FsAverage template, a surface-based model of the cortical surface based on 40 MRI scans of real brains (11), the only source space implemented in the MNE toolbox. It comprises 5124 solution points (equivalent to voxels in fMRI) equidistantly spaced throughout the surface of the cortical gray matter including insular cortex. We used a realistic three-layer (brain, skull, scalp) boundary-element model to construct the lead-field and solve the forward model. The so-created inverse operator (dSPM (12)) was then applied time-pointwise to the evoked responses averaged

across all trials in each subject and each condition to reconstruct the intracranial current density at each of the 5124 solution points.
Clinical and experimental studies consistently demonstrate that carefully conducted source reconstructions yield reliable estimates of intracranial currents. To avoid spatial aliasing and blurring of focal sources by the impedance of the skull and scalp, it is crucial to record from a sufficiently high number of electrodes, with 64 considered the minimum for reliable source estimations (13). The gold-standard of reliability estimates of intracranial source reconstructions are studies in epileptic patients that directly compare source-reconstructions from scalp-recordings of interictal epileptic spikes with intracranial and peri-operative recordings and fMRI in the same patients that are post-operatively seizure-free, i.e. in whom the focal source of interictal activity could be correctly localized from scalp recordings (13–25). At this point, it is important to note that interictal activity is often non-cortical but generated in deeper medial temporal areas, which can be as reliably reconstructed as more superficial sources. One can argue that interictal spike activity has sharper peaks and larger amplitudes and thus an inherently large signal-to-noise ratio and is hence more easily reconstructed. However, there is ample evidence from experimental studies that identified the same hippocampal sources reconstructed from surface EEG in healthy subjects (26) and in intracranial recordings in epileptic patients (27) in the same experimental paradigm.
Many studies in healthy human subjects have yielded sensible and reliable source estimates of cortical and deeper sources in a wide range of experimental paradigms ranging from the correct localization of auditory (28), visual and somatosensory (29) and even olfactory evoked potentials (30) and a variety of cognitive experimental paradigms using both visual and auditory stimuli (31–39) including functional localizers comparable to those used in fMRI (40) and by direct comparison of EEG source localization and fMRI in simultaneous recordings (41) and recordings in the same subjects (42).

### Identification of intracranial generator differences

ERP amplitude differences measured as the scalp surface can arise from differences in 1) strength or 2) timing of the same generator or 3) from different generators (that have different configurations).
Reconstructing sources from ERP difference waves is problematic for multiple reasons: difference waves are obtained by subtracting the ERPs in one condition from another, and the direction of subtraction is inherently arbitrary (A-B = -(B-A) with |A-B| = |B-A|); moreover, if amplitude differences arise from different topographies, the amplitude difference wave does not adequately reflect the spatial summation of the underlying generators in different experimental conditions. More importantly, intracranial current density estimates [in $mA/mm^3$] are an unsigned measure: they can assume only positive values of different magnitudes (i.e., there is current density of a certain magnitude or there is nearly none, but there cannot be “negative” current density), and because they are inherently “blind” to the direction of an effect obtained from a difference wave, they yield comparable estimates for A-B and B-A; see (36) for an in-depth discussion of this

issue. The most straightforward way to overcome these inherent problems in localizing intracranial source differences between conditions is to apply time-point-and-voxel-wise statistical parametric mapping of intracranial source differences in the inverse space like in fMRI (31, 32, 35, 40). Linear contrasts (e.g., T-tests) comparing intracranial current density values between experimental conditions yield a signed value that reliably indicates in which condition the current density is larger. This also overcomes the problem of thresholding, i.e., above which a source is considered "active" and so eliminates the effects of spurious sources (due to measurement errors) that do not vary systematically between conditions. There is no and there cannot be an a-priori value above which a source should be considered as active: because the strength of the electrical field declines with the inverse of the squared distance, different values should be assigned to superficial cortical in gyri and sulci and deep sub-cortical sources. Thresholding by means of statical comparisons avoids any bias in determining when a source should be considered as active. Numerous experimental studies applying source space statistics have yielded reasonable source estimations (26, 31–40, 42).

The multiple comparison problem in fMRI also applies to source space statistics because of the inherently large number of comparisons (solution points (5124) x time points). Traditional approaches which either control the familywise error rate (e.g., the Bonferroni correction) or the false discovery rate (FDR correction) do not take into consideration the highly correlated structure of the EEG data, especially in the source space. This leads to excessively conservative estimates prone to type II errors because the significance level is divided by the sum of all tests (solution points x time points). This problem can be addressed by clustering spatially and temporally coherent T-values above a pre-determined threshold ($p < .05$) to capture the spatio-temporal structure of the data (43, 44). The cluster statistics is calculated by taking the sum of all T-values that belong to the cluster. Statistical significance is then assessed creating a null distribution of cluster sizes using non-parametric permutations by randomly shuffling condition labels 5000 times, (this label shuffling preserves the inherent spatio-temporal structure of the data). To control for type I errors, only the sum of T-values belonging to the largest cluster is retained for each permutation. The cluster statistics is then tested against the null distribution of maximal T-values: the p-value of the cluster obtained from the experimental data is calculated as the proportion of maximal clusters that are larger than the original cluster statistics. If this proportion is less than 5%, it is concluded that the cluster size significantly deviates from the null. Drawing the null distribution directly from the clustered data preserves the spatio-temporal correlation structure in the data. "Significance" for every solution point is expressed in the temporal extend (in ms) of its contribution to the significance across the extend of the cluster (rather than in terms of statistical parameters). This sacrifices spatio-temporal *precision* for spatio-temporal *consistency* across the cluster, in other words, it indicates how consistently each point contributes to the spatio-temporal extent of activity differences in the entire cluster.

Figure S1

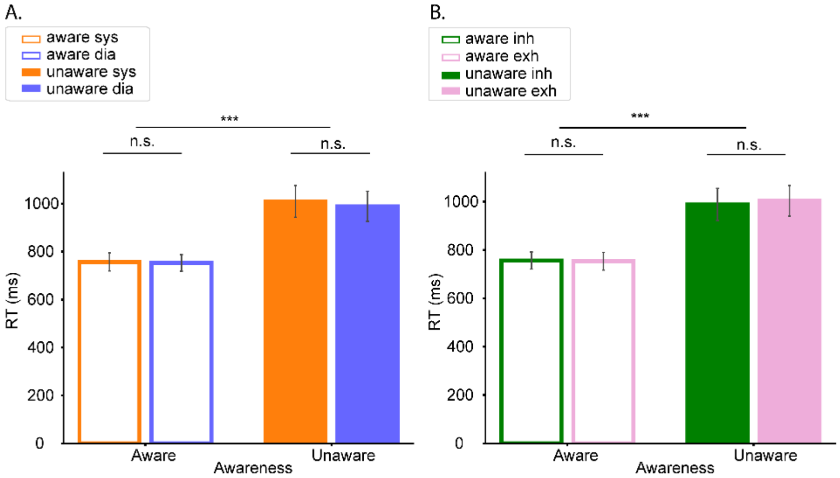


**Fig. S1.** Behavioral results: average reaction times (in ms) as a function of awareness and (A) cardiac phase (systole: orange, diastole: blue) and (B) respiratory phase (inhalation: green, exhalation (pink)). The error bars represent the standard error, *** p<.001.

Figure S2

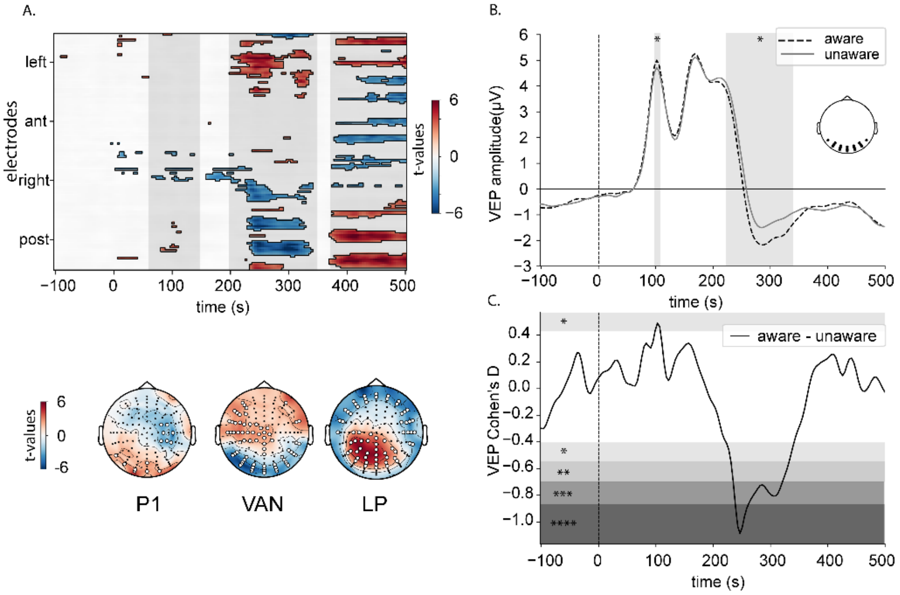


**Fig. S2.** Main effect of awareness on ERP amplitudes. (A) Time-course and location of significant differences for the contrast between aware and unaware identification; top panel: t-values denoting the direction of the main effect (identified by mass univariate ANOVAs, FDR corrected across time and space)); three periods of significant amplitude differences are highlighted in gray: P1 (90 – 120 ms) VAN (250 – 350 ms), P3/LPC (400 – 500 ms), bottom panel: topo-maps illustrating the distribution of the main effects; electrodes showing a significant electrodes main effect of awareness are highlighted by a white circle. (B) Grand average ERP waveforms averaged across the electrodes showing a main effect of awareness in the time-window of the VAN (250 – 350ms) displayed in the inset (C) Time-course of Cohen's d effect sizes for the contrast aware – unaware at the electrodes highlighted in the inset in panel (B); gray shades indicate significance levels (*p<0.05, * p<0.01, ***p<0.001, ****p< 0.0001).

Figure S3

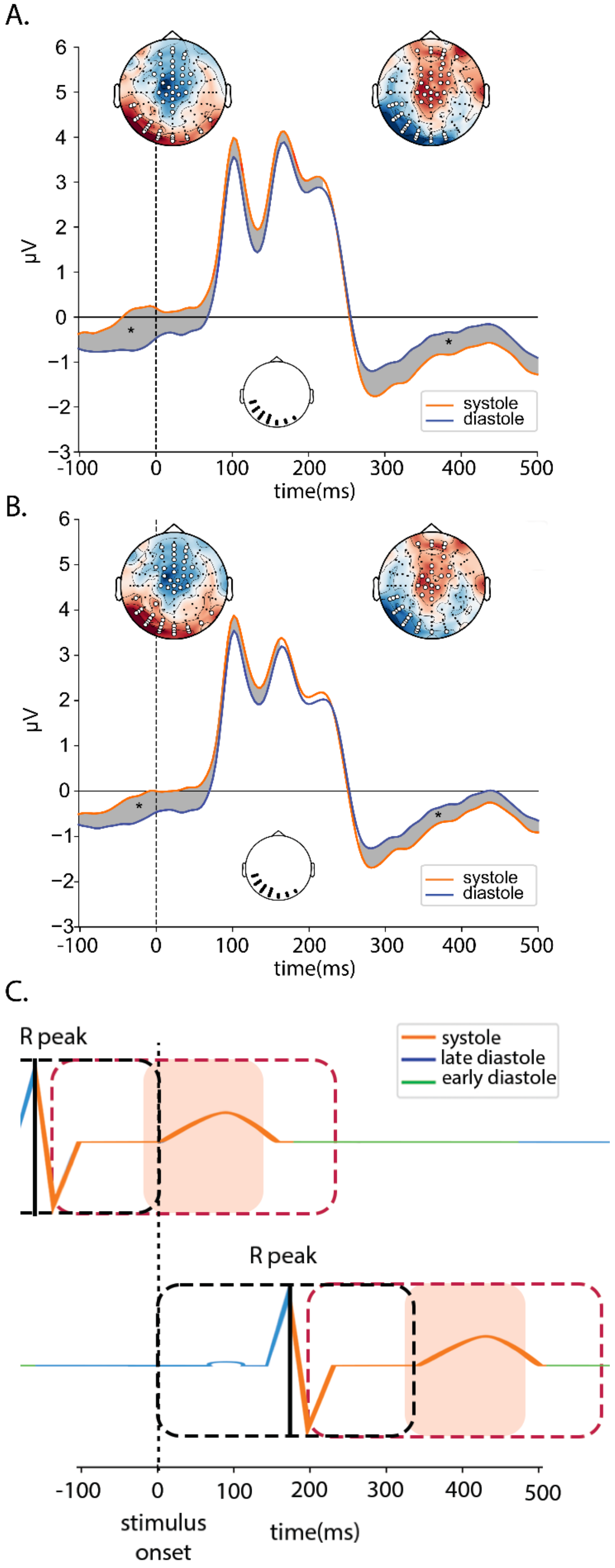


**Fig. S3.** Main effect of cardiac phase on ERPs. (A) Grand Average ERP waveforms averaged across the occipito-temporal electrodes highlighted in the insert in the systole (orange) and diastole (blue) without correction of the cardiac field artifact (CFA). Gray shades indicate time-periods of a significant main effect of cardiac phase identified by mass univariate ANOVAs (FDR corrected), topoplots illustrate the distribution of the main effect, electrodes showing a significant main effect of awareness are highlighted by a white circle. (B) Grand Average ERP waveforms averaged across the occipito-temporal electrodes highlighted in the insert in the systole (orange) and diastole (blue) with correction of the CFA. (C) Illustration of the time-course of the cardiac pulse wave relative to stimulus onset. The dotted black line denotes stimulus onset; the dashed black and red windows denote the time window in which the R peaks and the ensuing pulse waves can occur for stimuli falling into the systole (top panel) and diastole (bottom panel); the solid black line and the orange highlight indicate an exemplar R peak and pulse wave in each condition; note how the significant main effect of cardiac phase in panels (A) and (B) align with the red dashed windows in panel (C).

Figure S4

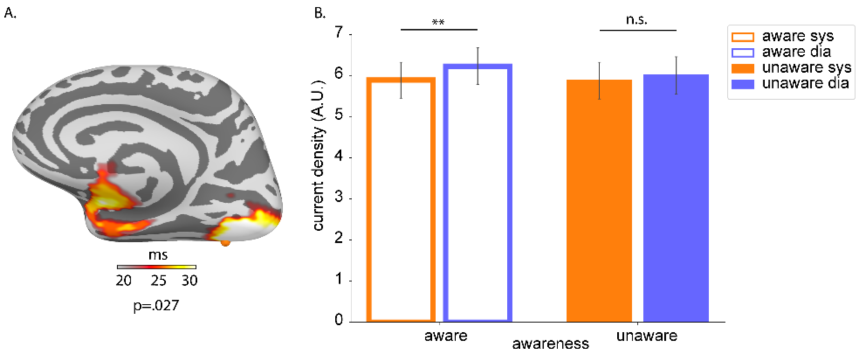


**Fig S4**. Current density differences in extrastriate cortex underlying the interaction between awareness and cardiac phase during the P1. (A) Statistically significant difference in current density in the aware condition between diastole and systole rendered on the FsAverage template; the color bar indicates the duration of significance (> 20ms at p=0.027) for each point in the cluster; the orange dot indicates the location of maximal activation duration (x=-34.3, y=-73.5, z=-12 mm). (B) Current density averaged across all points in the cluster displayed in (A) in the systole (orange) and diastole (blue) as a function of awareness; error bars display the standard error; **p<0.01

Figure S5

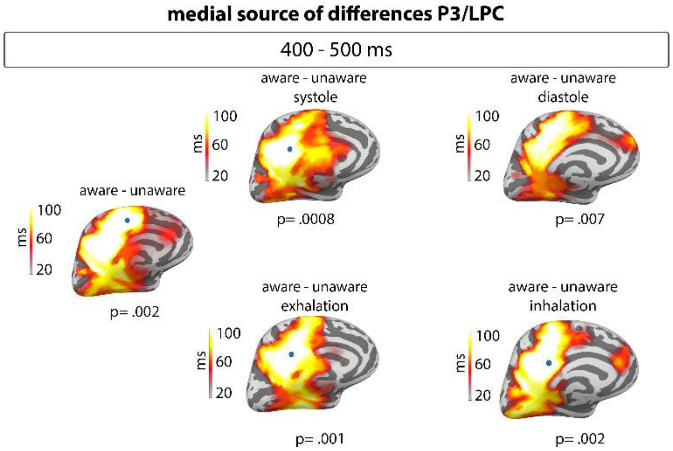


Fig S5. Medial extent of current density differences underlying the main effect of awareness as a function of the cardiac and respiratory phase for the P3/LPC component rendered on the FsAverage surface template; color bars indicate the duration (>20 ms) of significant activity in the spatio-temporal clusters between the aware and unaware conditions at the respective significance level. Left column: main effect of awareness irrespective of cardiac or respiratory phase. Top row: main effect of awareness irrespective in the systole (left) and diastole (right). Bottom row: main effect of awareness during exhalation (left) and inhalation (right).
